# Single calcium channel nanodomains drive presynaptic calcium entry at lamprey reticulospinal presynaptic terminals

**DOI:** 10.1101/2021.02.02.429388

**Authors:** S Ramachandran, S Rodgriguez, M Potcoava, S Alford

## Abstract

Synchronous neurotransmission is central to efficient information transfer in neural circuits, requiring precise coupling between action potentials, Ca^2+^ entry and neurotransmitter release. However, determinations of Ca^2+^ requirements for release, which may originate from entry through single voltage-gated Ca^2+^ channels, remain largely unexplored in simple active zone synapses common in the nervous system. Understanding these requirements is key to addressing Ca^2+^ channel and synaptic dysfunction underlying numerous neurological and neuropsychiatric disorders. Here, we present single channel analysis of evoked active zone Ca^2+^ entry, using cell-attached patch clamp and lattice light sheet microscopy over active zones at single central lamprey reticulospinal presynaptic terminals. Our findings show a small pool (mean of 23) of Ca^2+^ channels at each terminal, comprising subtypes N-type (CaV2.2), P/Q-type (CaV2.1) and R-type (CaV2.3), available to gate neurotransmitter release. Significantly, of this pool only 1-6 (mean of 4) channels open upon depolarization. High temporal fidelity lattice light sheet imaging reveals AP-evoked Ca^2+^ transients exhibiting quantal amplitude variations between action potentials and stochastic variation of precise locations of Ca^2+^ entry within the active zone. Further, Ca^2+^ channel numbers at each active zone correlate to the number of presynaptic primed synaptic vesicles. Together, our findings indicate 1:1 association of Ca^2+^ channels with primed vesicles, suggesting Ca^2+^ entry via as few as one channel may trigger neurotransmitter release.

## Introduction

Presynaptic action potentials evoke Ca^2+^ entry through voltage-gated Ca^2+^ channels (VGCCs) in the presynaptic terminal.^1^ However, with the exception of specialized calyceal synapses^2,3^, it remains unclear how much Ca^2+^ is required to trigger vesicle fusion and neurotransmitter release and, in lieu of this, how many channels open upon depolarization. It is clear, that as vesicles dock and prime at presynaptic active zones (AZ), VGCCs tether or are tethered to the fusion machinery.^4–6^ Precise estimates of Ca^2+^ requirements for release, and the relationships between VGCCs and their targets at the fusion machinery, is vitally important to understand how synaptic transmission occurs and how it is subject to modulation and plasticity.^7^ Here, we present single channel analysis of VGCCs at single lamprey reticulospinal AZs by cell-attached patch clamp electrophysiology. We confirm these findings by performing high-fidelity imaging of fluorescent Ca^2+^ transients, using a unique lattice light sheet microscope (LLSM) to image single action potential (AP)-evoked fluorescent Ca^2+^ transients *in situ*. Our findings indicate presynaptic Ca^2+^ entry at these synapses occurs through few open channels, possibly even one, localized in close proximity to the synaptic vesicle at the release site, and present a novel advancement towards understanding Ca^2+^ regulation of release at central synapses.

## Results

### Dissociated axons retain functional terminals

Precise measures of evoked presynaptic Ca^2+^ influx requires direct recordings from the release face membrane of individual presynaptic terminals, impeded *in vivo* at central synapses due to the close apposition between the presynapse and postsynapse [Fig.1A]. Such recordings have, thus far, been possible only at specialized calyceal-type terminals.^2,3^ We developed an acute dissociation of the lamprey spinal cord^8^ [Fig.1B, Methods], enabling isolation of reticulospinal axons with functional presynaptic terminals without any postsynaptic processes [Fig.1B-C]. Dissociated axons retained structural integrity [Fig.S1A], physiological membrane potentials (−58 ± 1.6 mV, n=17 axons) and the capacity to fire APs (n=7 axons) [Fig.S1B]. Most notably, presynaptic terminals maintained evoked synaptic vesicle exo-endocytosis visualized by FM1-43 staining of recycling vesicle clusters during 30 mM KCl depolarization [Fig.1C,S1C],^9^ enabling targeted recordings of Ca^2+^ currents from fluorescently identified single terminals [Fig.1C-D]. Notably, we recorded only from presynaptic terminals with simple AZs (diameter 0.4 – 2.8 µm, mean 1.6 ± 0.2 µm), with single release sites and univesicular release,^10–12^ allowing for correlations of Ca^2+^ measures presented in this work to the fusion requirements of a single synaptic vesicle.

**Figure 1:**
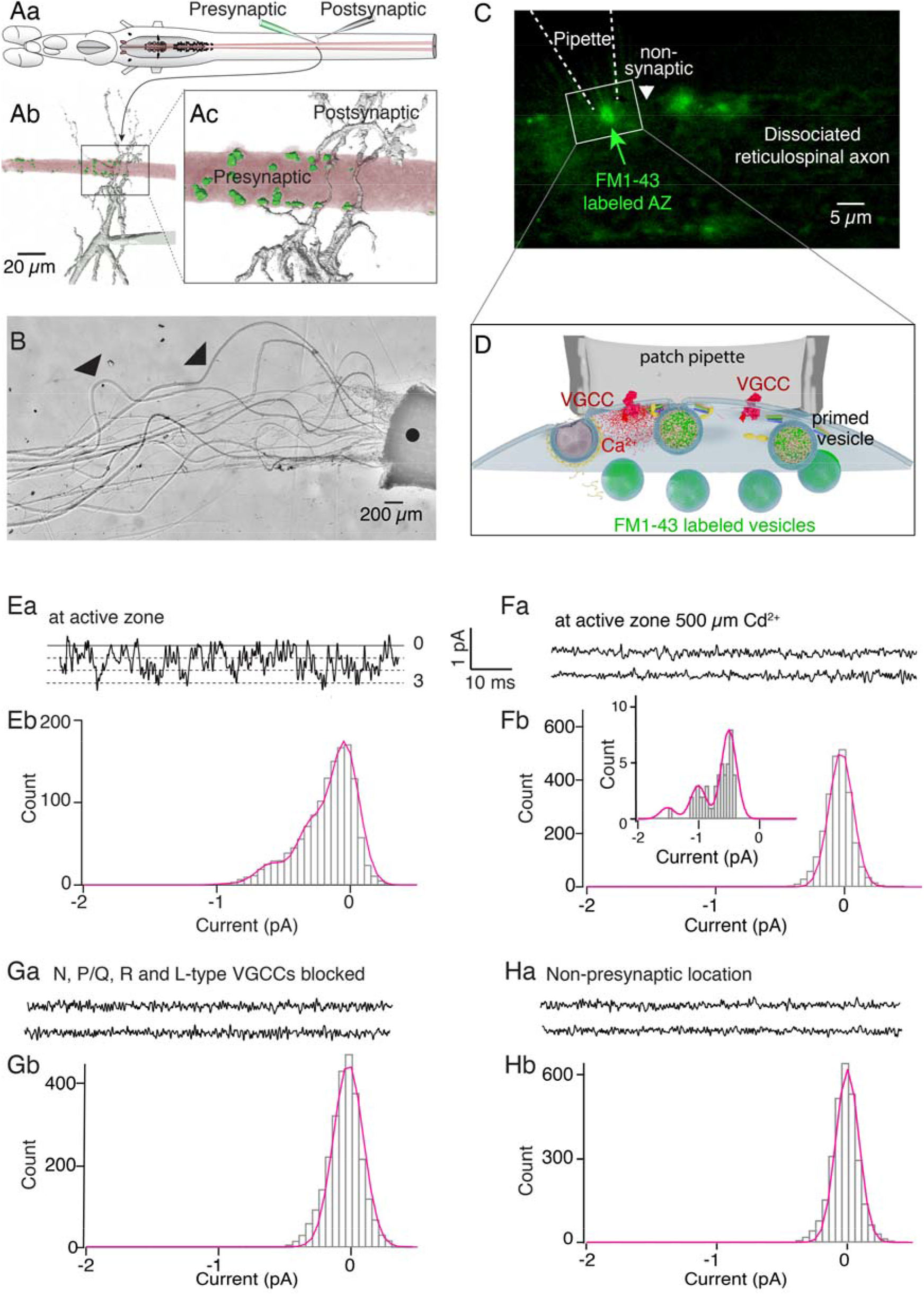
Direct recording of VGCC currents at the release face membrane of individual AZs. **A**. [Aa] Schematic of the lamprey spinal cord showing reticulospinal axon/spinal neuron synapses demonstrating paired recording. [Ab] Image depicting the architecture of the reticulospinal synapse. Presynaptic axon is colored pink, presynaptic terminals are in green (from intracellular phalloidin labeling of actin surrounding vesicle clusters in the axon^9,46^). Postsynaptic projections (grey) contact presynaptic terminals. [Ac] Magnified image of black inset box in Ab. Scale bar 20 µm. **B**. Giant axons were acutely dissociated. Representative image showing the dissociated reticulospinal axon preparation. Solid black circle indicates dissociated spinal cord end from which reticulospinal axons emerge out. Arrowheads (black) indicate reticulospinal axons with no apposing postsynaptic processes. Scale bar 200 µm. **C**. Representative image of cell attached patch clamp recording at a single presynaptic AZ of a dissociated reticulospinal axon labeled with FM 1-43 (green puncta). Dashed white lines outline the pipette position. White arrowhead indicates non-presynaptic terminal location. Scale bar 5 µm. **D**. Schematic showing placement of a cell attached electrode over the AZ identified by FM 1-43 labeling of recycling vesicles. These are present in a defined cluster immediately adjacent to the active zone colocalized with actin labeled with phalloidin [Fig.1Ab-c]. The vesicle cluster margins extend beyond the active zone^46^ indicating that the patch electrode covers the entire structure. **E**. [Ea] Representative sweep from a cell attached patch recording showing Ca^2+^ currents recorded from a single AZ following a voltage step to 0 mV (n=5 patches). For all Ca^2+^ current traces shown, inward currents have a negative directionality. Solid line marked 0 indicates channel closed state, dashed lines indicate incremental channel opening events corresponding to the opening of 1, 2 or 3 channels. Patch pipette contains 10 mM [Ca^2+^]_external_ as charge carrier. [Eb] Amplitude histogram (grey bars) fitted by a sum of 3 Gaussian function (magenta fit line), indicates the opening of 1-3 Ca^2+^ channels during the indicated recording. **F**. [Fa] Similar cell-attached recordings, from single AZs, following a voltage step to 0 mV with 500 µM Cd^2+^ and 10 mM [Ca^2+^]_external_ in the patch pipette demonstrating absence of inward Ca^2+^ currents. [Fb] Amplitude histogram (grey bars) (n=8 patches) fitted with a single Gaussian function (magenta fit line) indicating only a current noise peak and no channel events. [Inset] Tail currents measured, upon repolarization to −80 mV from the voltage step during release of Cd^2+^ block, plotted as amplitude histogram (grey bars). The amplitude histogram was fitted by a sum of 3 Gaussian function (magenta fit line), indicating the opening of 1-3 Ca^2+^ channels upon repolarization. **G**. [Ga] Cell-attached recordings from single AZs, following a voltage step to 0 mV, showing absence of depolarization evoked inward currents when all VGCC subtypes (N, P/Q, R and L) were blocked. [Gb] Amplitude histogram (grey bars, n = 4) fitted with a single Gaussian function (magenta fit line) indicating only a current noise peak and no channel events. **H**. [Ha] Cell-attached recordings from non-presynaptic terminal locations, [white arrowhead in Fig.1C], following a voltage step to 0 mV, demonstrating absence of inward Ca^2+^ currents. [Hb] Amplitude histogram (grey bars, n = 3) fitted with a single Gaussian function (magenta fit line) indicating only a noise peak and no channel events.

### Multiple VGCC subtypes at reticulospinal AZs

We validated that VGCC localization to the active zone release face membrane was retained post dissociation, using antibodies specifically labeling three different Ca^2+^ channel subtypes, N-type (CaV2.2), P/Q-type (CaV2.1) and R-type (CaV2.3) [Fig.S2-3], previously implicated in synaptic transmission from lamprey reticulospinal axons^9,13,14^.s

We recorded Ca^2+^ currents at the release face membrane of individual fluorescently labeled AZs [10 mM [Ca^2+^]_external_ (in patch pipette), Fig.1E, n=5 patches] using custom low-noise single channel cell-attached patch clamp recordings^8^. Ca^2+^ currents were evoked by applying a depolarization step protocol [Fig.2A] from holding potential (−80 mV – adjusted from the known intracellular membrane potential determined from intracellular sharp electrode recording). We isolated VGCC subtype-specific currents pharmacologically [Methods], with 10 mM [Ca^2+^]_external_ (in patch pipette), eliciting Ca^2+^ currents corresponding to four different VGCC subtypes: N-type (CaV2.2, n=6 patches), P/Q-type (CaV2.1, n=5 patches), R-type (CaV2.3, n=6 patches) and L-type (CaV1, n=4 patches) [Fig.2B]. Channel openings were observed only upon depolarization to a membrane voltage of −30 mV [Fig.2B,3B], indicating the absence of any low-voltage-activated T-type (CaV3) channels. Pharmacological block of all VGCCs with Cd^2+^ [Fig.1F, n=8 patches] or with combined selective blockers of all four VGCC subtypes [Fig.1G, n=5 patches] eliminated all currents. Additionally, no currents were recorded when patch recordings were obtained from non-presynaptic terminal locations determined by the absence of fluorescent FM 1-43 puncta along the axon membrane [Fig.1H, n=3 patches]. We measured single channel amplitude (i), for each channel subtype from current records that clearly demonstrated the opening of only a single channel, at depolarized voltages of −30 mV, 0 mV, 30 mV and upon repolarization to −80 mV (tail currents) [Fig.2A, Supplement]. Single channel amplitudes at 0 mV were 0.28 ± 0.01 (N-type), 0.29 ± 0.03 (P/Q-type), 0.28 ± 0.02 (R-type) and 0.28 ± 0.02 pA (L-type) [Fig.2C, Table.S1]. Single channel conductance was calculated to be 2.97 ± 0.62 pS (N-type), 1.82 ± 0.36 pS (P/Q-type), 1.94 ± 0.41 pS (R-type) and 2.4 ± 0.75 pS (L-type) [Fig.2C, Table.S1]. Mean channel opening probability (P_open_) at 0 mV was 0.26 ± 0.02 (N-type), 0.29 ± 0.03 (P/Q-type), 0.22 ± 0.02 (R-type) and 0.24 ± 0.02 (L-type) [Fig.2D, Table.S2].

**Figure 2:**
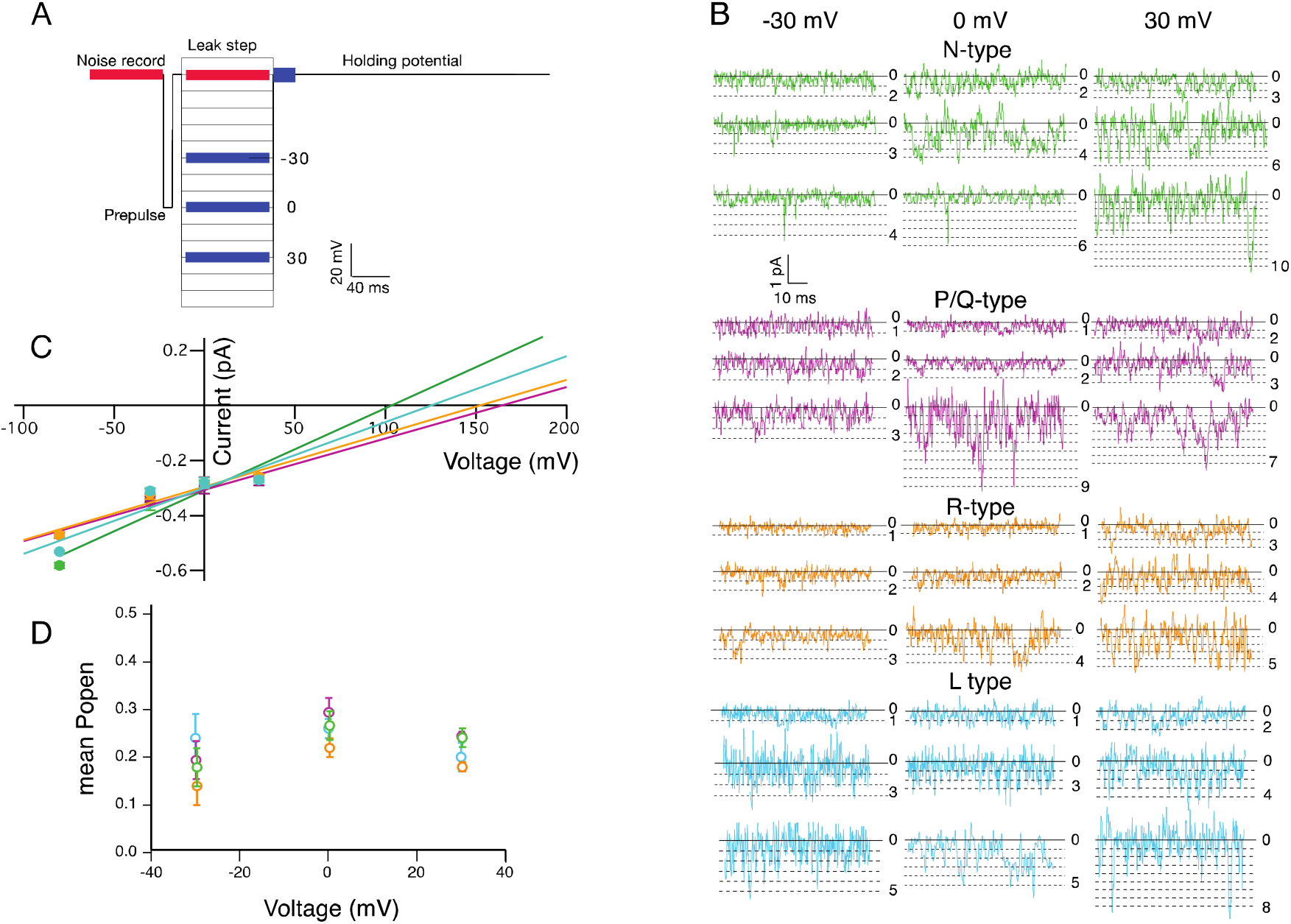
Cell attached recordings from AZs reveal multiple VGCC subtypes. **A**. Depolarization stimulus protocol, applied from a holding potential of −80 mV, to elicit VGCC currents in cell-attached patch recordings. Red bar at the holding potential level indicates where the background current noise was measured (current noise record). Blue bars indicate the voltages at which analysis of single channel currents (current records) was carried out: step depolarizations to −30 mV, 0 mV and 30 mV, and upon repolarization to −80 mV (tail currents). **B**. Representative examples of N-type (CaV2.2), P/Q-type (CaV2.1), R-type (CaV2.3) and L-type (CaV1) currents, recorded at single AZs, following voltage steps to −30 mV, 0 mV and 30 mV. Solid line marked 0 indicates channel closed state, dashed lines indicate incremental channel opening events, with maximum number of simultaneously open channels indicated in each representative trace. **C**. Current-Voltage relationships plotted for each of the indicated Ca^2+^ channel subtypes. For each channel subtype, mean single channel amplitude measured at corresponding voltages is shown along with the linear fit (N-type green, P/Q-type purple, R-type orange, L-type blue). Error bars indicate SEM. Linear fits have been extrapolated to intersect the X-axis indicating the reversal potential (E_rev_). **D**. Mean open probability (mean P_open_) plotted against membrane voltage (mV) for each of the indicated Ca^2+^ channel subtypes (N-type green, P/Q-type purple, R-type orange, L-type blue).

### Few open channels from a small available VGCC pool mediate Ca^2+^ influx

We next enumerated Ca^2+^ channels at single presynaptic AZs for each Ca^2+^ channel subtype. The current data from replicate sweeps, for each analyzed voltage, in a patch recording were binned to generate amplitude histograms, which were averaged and fitted with a sum of Gaussians function [Supplement]. Using this approach, we enumerated the maximum number of channels, visually confirmed by inspection of the channel opening events, that simultaneously opened at each analyzed voltage during a patch recording. A maximum count of 10 (4-10, mean 5) N-type, 9 (3-9, mean of 6) P/Q-type, 32 (4–32, mean 12) R-type and 17 (3–17, mean 10) L-type channels, yielded a total maximum VGCC count of 68 channels (mean 33) [Fig.3A, Table.S3] at a presynaptic AZ. Restricting the count to only N+P/Q+R channel subtypes, that are implicated in evoked synaptic transmission at lamprey reticulospinal synapses,^9,13,14^ we obtained a maximum total of 51 channels (mean 23) [Fig.3A, Table.S4] that may contribute Ca^2+^ to drive release at a single AZ. Having determined the available pool of VGCCs at a single AZ, we next asked how many VGCCs open upon depolarization, when all channel subtypes are available for opening. We carried out cell-attached recordings from individual presynaptic AZs [Fig.1C-D], with 90 mM [Ba^2+^]_external_ (in patch pipette). Upon depolarization to 30 mV, we observed 1-6 (mean 4) open channels at an AZ [Fig.3B, Table.S5, n=5 patches], suggesting very few out of the available small pool of channels mediate Ca^2+^ influx at depolarization voltages that are equivalent to peaks of action potentials.

**Figure 3:**
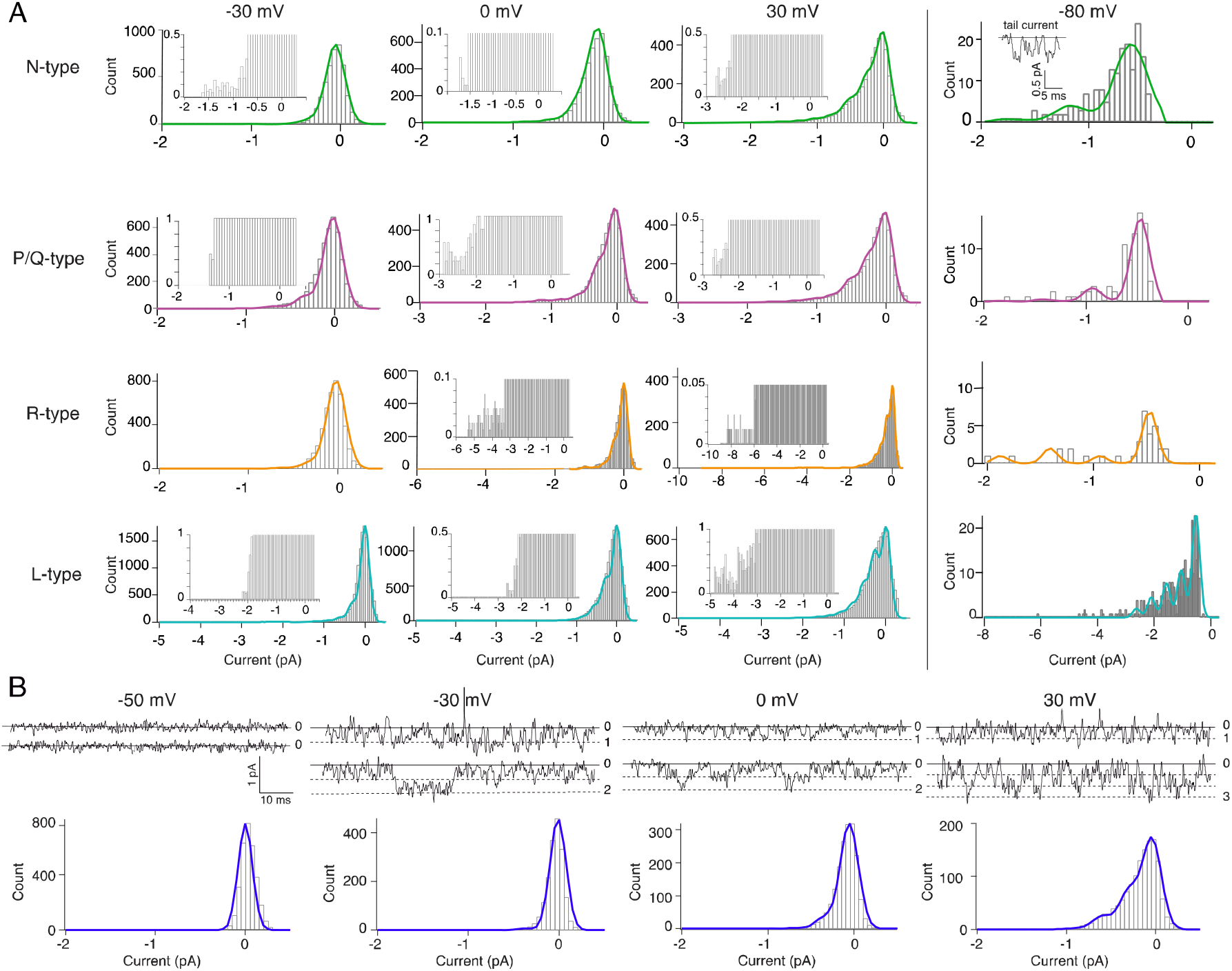
Few open channels from a small available VGCC pool mediate Ca^2+^ influx. **A**. Gaussian fits for the amplitude histograms (grey bars) for the indicated Ca^2+^ channel subtypes (10 mM [Ca^2+^]_external_) following voltage steps to −30 mV, 0 mV and 30 mV. Tail currents were measured, within a window of 20 ms immediately after repolarization to −80 mV, and plotted as amplitude histograms (grey bars). Inset (in N-type −80 mV) shows representative example of tail current channel opening events. Sum of Gaussian fit to the amplitude histogram is shown as solid line (N-type green, P/Q-type purple, R-type orange, L-type blue). Insets in histogram plots provides magnified view of the histogram (note expanded Y axis) showing bin values for higher current amplitudes that are not discernable from the full scaled histogram. **B**. Representative current traces, from cell attached patch clamp recordings at single AZs with 90 mM [Ba^2+^]_external_ in the patch pipette, following voltage steps to −50, −30, 0 and 30 mV. Solid line in all the representative traces indicates the closed state of the channel. Dashed lines indicate incremental channel opening events, with maximum number of simultaneously open channels indicated in each representative trace. Amplitude histograms are shown in grey and the sum of Gaussian fit line in dark blue. Note the absence of any channel openings at a membrane voltage of −50 mV, 1-2 channels at −30 mV and 1-3 channels at 0 mV and 30 mV.

### Molar quantitation corroborates Ca^2+^ entry through few open VGCCs

To validate our single channel recording findings, we characterized stimulus-evoked Ca^2+^ influx into the presynaptic terminal *in situ*, by introducing exogenous fluorescent Ca^2+^ dyes as buffers (Fura-2, Oregon Green BAPTA1 and Fluo5F) with different affinities into the terminal at known concentrations. We recorded evoked Ca^2+^ entry upon repetitive AP firing [Fig.S4A].^15,16^ Knowing κ_dye_ (buffering capacity of the exogenous dye) within the axons during stimulation, the endogenous buffering capacity of the axons κ_endo_ was estimated and action potential-evoked free Ca^2+^ concentrations and total concentrations of Ca^2+^ entering terminals were determined [Fig.S4B-D, Table.S6, Supplement]. We also determined the total axonal volume that this Ca^2+^ entered into and the number of active zones responsible. From this, we calculated the total molar amount of Ca^2+^ entering a single active zone to be 4.14 ± 0.47 nMoles /stimulus, yielding a charge of 7.98 ± 0.90 fC or a sustained current of 1.60 ± 0.22 pA over the AP duration (5 ms – including tail current duration of 2 ms). This corresponds to the opening of 3.6 to 4.8 channels when mean channel currents are calculated from the integrals of line fits [Fig.S4C] over the voltage range of the AP (−80 to 20 mV). These measures agree closely with the peak numbers of channels opening in direct cell-attached recordings [Fig.3B, Table.S5].

### Quantal variation in presynaptic evoked Ca^2+^ transients

Few channels opening on presynaptic depolarization suggest a very low coupling ratio between channels and primed vesicles, and implies that channel activation contributes to the stochastic nature of quantal release. We visualized effects of channel opening at single presynaptic terminals by imaging fluorescent Ca^2+^ transients using a customized LLSM^17^ [Fig.4A]. Lamprey giant synapses are uniquely suited for this study, because they have simple active zones^10^ and a large axon into which Ca^2+^ can disperse^18^ enabling resolution of Ca^2+^ transients from single active zones. LLSM provides a very narrow excitation thickness in z (∼0.4 µm), very high signal to noise and nearly no photobleaching.^17^ We imaged AP-evoked Ca^2+^ transients at AZs *in situ* in axons pre-loaded with an intermediate affinity Ca^2+^ dye (Fluo 5F, K_d_ 2 µM at 10°C) at high frame rates (330 to 800 Hz) over many repetitions at fixed z-axis [Fig.4B].

**Figure 4:**
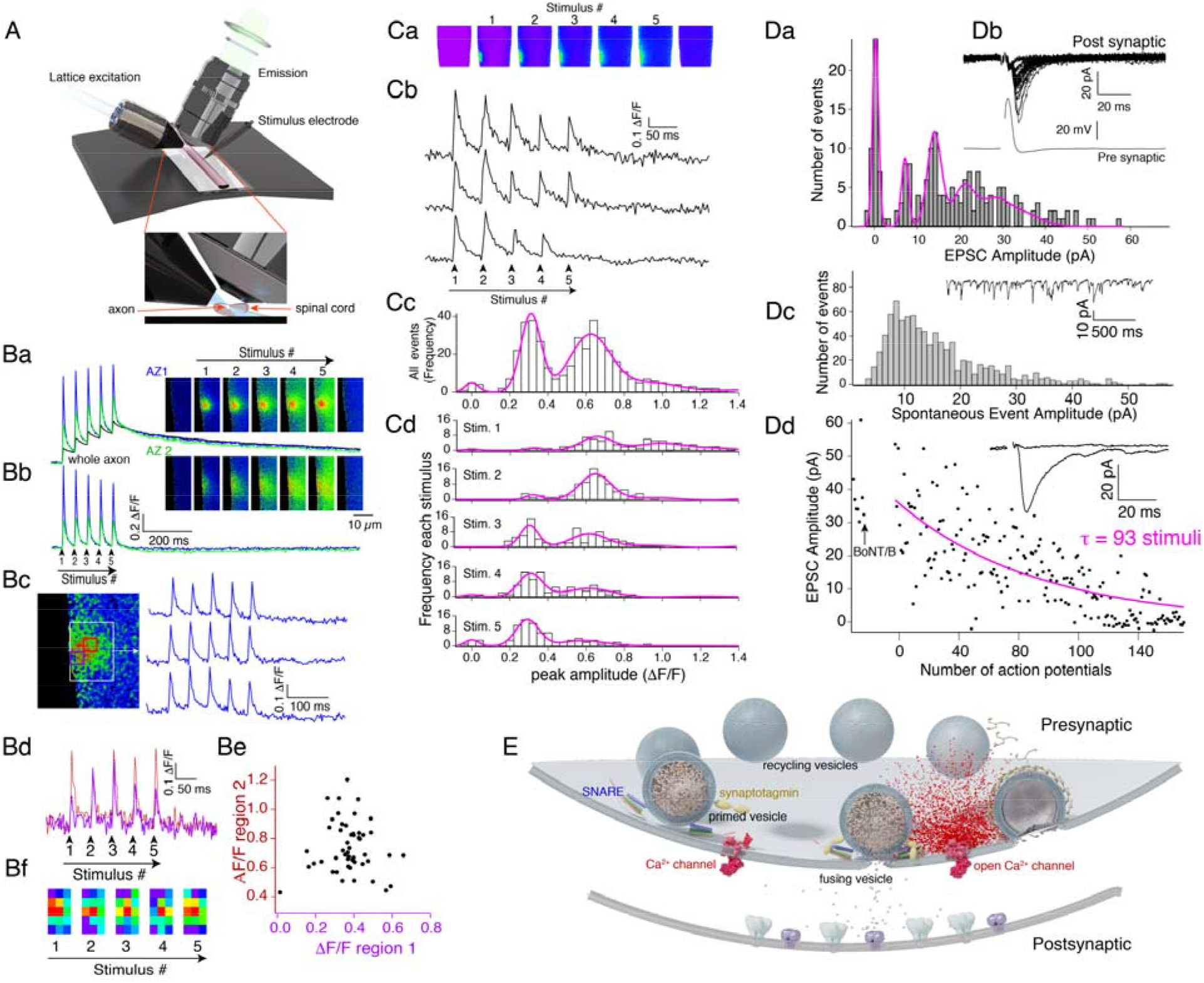
The relationship between channel numbers and vesicles at Azs. LLSM enabled electrophysiology and imaging at an active zone during stimulation in the intact preparation. LLS illumination (30 bessel beam, 51 × 51 µm) is projected onto a preparation pinned to sylgard. Orthogonal to this light sheet emission was imaged through a 1.1NA 25x lens to generate 2D data frames during stimulation of axons in intact spinal cord. (57 AZs, 18 preparations) **A**. [Ba] Trains (5 APs, 20 Hz) were stimulated with an electrode over ventro-medial tracts at axons pre-filled with Ca^2+^-sensitive dye (Fluo5F) and transients resolved at 300 Hz. [Bb] Two resultant hotspots are shown expressed as (ΔF/F; AZ1 blue, AZ2 green). To isolate local Ca^2+^ signals at single AZs the bulk axonal signal was subtracted from the AZ signal. [Bc] Amplitudes recorded from single AZs (ROI shown as white box) varied between stimuli. Sub-regions within this ROI showed larger variance. [Bd] two such subregions were further analyzed (red and purple boxes) shown as red and purple traces. [Be] Graph plots amplitudes from the 2 regions in [Bd] for 50 action potentials showing little correspondence between these sub-regions of the AZ. [Bf] a color look up coding of peak amplitudes for a 3 × 5 array of subregions from 1 train (each 1 µm^2^) from white boxed ROI in [Bc] showing poor covariance between regions and APs. **B**. Variation in Ca^2+^ transients between action potentials. [Ca] Single AZs were imaged (up to 320 action potentials in 80 trains) with no bleaching. [Cb] Transient hotspots were recorded as for [Bb]. Individual whole AZ transients were measured to give peak amplitudes. Plotting all peaks as a histogram for all APs [Cc] shows distinct peaks implying events are multiples of a unit amplitude. Gaussian fits (magenta) were made to this distribution with two constraints (1) amplitudes between peaks were equal, (2) variance scaled linearly with increase in amplitude. A similar set of histograms were calculated for each of the first through the fifth stimulus and gaussians were fit. While the numbers of low amplitude responses increased later in the train the unit amplitudes remained constant. **C**. Counting primed vesicles. [Da] P_r_ was calculated from the quantal distribution of event sizes from paired cell recording (n=4), whereby axons were stimulated through a microelectrode and postsynaptic currents recorded in whole cell (inset: recordings from 30 stimuli). Histograms of 200 sequential stimuli were fit with multiple gaussians (magenta) [Db]. Spontaneous event amplitudes we determined in further paired recordings. La^3+^ was injected into presynaptic axons (n=4) to increase spontaneous event frequency from the recorded axon. Amplitude distribution histograms were constructed from these data. [Dc] To determine the number of APs necessary to deplete the primed vesicle pool, further paired recordings were made. [Dd] BoNT/B was injected into the presynaptic axon. After 5 mins to allow toxin to act, stimuli (1Hz) were applied and the EPSC measured until it decayed to zero. An exponential fit (magenta) to this gives a decay (τ) of 93 stimuli. **D**. Schematic model of the terminal. A small number of primed vesicles are associated with an equal and equally small number of Ca^2+^ channels available at the presynaptic terminal. Of these channels only (0-6) open in one active zone on each action potential. This provides part of the reason for low probability of release.

Superficially, Ca^2+^ transients appear as single hotspots [Fig.4Ba-b], that prior studies demonstrate colocalize to AZs.^9^ LLSM reveals a readily resolved transient during each AP [Fig.4Bc]. However, analysis of sub-regions within a single AP-evoked transient [AZ1, Fig.4Ba-b] shows substantial amplitude (peak ΔF/F) variance between APs within short trains and between trains [Fig.4Bd]. Peak amplitude varied stochastically at each sub-region, with different sub-region amplitudes not covarying at each AP [Fig.4Be]. We also analyzed Ca^2+^ transients, encompassing the whole AZ, to determine their amplitude variation between APs. Histograms of the amplitude distributions were well fit by 2-6 gaussians. [Fig.4Cc, 6 AZs] These may represent quantal amplitudes or individual channel openings, in addition to a peak at zero indicating channel opening failure. For all 6 axons tested, if each gaussian corresponds to a channel event, then a modal number of 1.8 ± 0.8 channels open at each AZ with a minimum of 0 and maximum of 7. Failures were not due to the absence of AP firing, because events were observed at other locations in the same axon on the same stimuli. In all AZs investigated, the mean amplitude of the response reduced from the first to the last event in the train. Analysis of amplitude distributions in each transient (1 to 5) over all trains [Fig.4Cd] reveals similar peak spacing but with higher probability of peaks at smaller values of ΔF/F and increased failures. These results are consistent with small numbers of channel openings with uniform amplitudes detected for AP and a reduction of opening probability at later stimuli in the train.

### 1:1 stoichiometry between Ca^2+^ entry through single open channels and single primed vesicles

We next related numbers of Ca^2+^ channels at AZs to numbers of primed vesicles at each AZ. In paired recordings between reticulospinal axons and spinal neurons, peak evoked EPSC amplitude histograms yielded a minimum evoked EPSC amplitude of 8.1 ± 2.7 pA [Fig.4Da-b, n=5]. We cannot directly compare this to a spontaneous event amplitude because there are too many non-reticulospinal synapses onto the postsynaptic cell.

However, asynchronous release was enhanced from just single reticulospinal presynaptic axons by injecting La^3+^ (1mM) to cause a tenfold increase in total spontaneous event frequency recorded in paired axons and neurons. This approach yielded a modal amplitude of asynchronous events (6.6 ± 1.5 pA, n=4), not significantly different from the minimal evoked amplitude (p = 0.58) [Fig.4Dc]. Mean probability of release [P_r_] was calculated from the gaussian fits [Fig.4Da] to be 0.22 ± 0.02. To understand the relationship between total numbers of VGCCs at AZs and numbers of primed vesicles, we calculated the number of APs required to deplete the primed vesicle pool [Fig.4Dd] by preventing vesicle priming with BoNT/B. This prevents neurotransmission by selectively cleaving synaptobrevin, but cannot cleave targets after formation of ternary SNARE complexes during priming^19^. In paired recordings, axons were injected with light chain BoNT/B^20^. Post BoNt/B injection, and after 5 mins to allow cleavage, the τ of EPSC amplitude decay expressed as number of APs, was 100 ± 17 [Fig.4Dd]. If we consider P_r_, the mean number of primed vesicles per AZ is 22, which corresponds very closely to the mean number of Ca^2+^ channels at the AZ (23), suggesting a possible 1:1 stoichiometry between AZ calcium channel numbers and individual primed vesicles. Together, our findings indicate an extreme nanodomain of calcium channels driving neurotransmitter release at these AZs, wherein a pool of available channels that corresponds in number to primed vesicles is available. APs evoke calcium entry through a subset of these VGCCs, possibly even a single open channel tightly coupled to the release site, to trigger vesicle fusion and release.

## Discussion

Rapid evoked neurotransmission is triggered by AP-coupled Ca^2+^ influx (< 60 µs)^21^, in close proximity to the fusion machinery^22^. However, how many VGCCs open to trigger fusion of a single vesicle remains unclear. While this has proven impossible to visualize up to now in simple low probability central synapses, direct recordings from larger and more specialized calyceal synapses^2,3^ reveal complex AZs with mixed properties. Single VGCCs (Ca^2+^ nanodomains) may gate release^23–27^, and in ciliary ganglion synapses shown to be tethered to docked synaptic vesicles^4^. In contrast, at other synapses multiple VGCCs (Ca^2+^ microdomains) drive release^28,29^, with a developmental shift to nanodomain-driven release at the calyx of Held^30,31^. Indirect recordings have suggested few VGCCs gate release in other synapses, such as frog neuromuscular junction^32^, granule cell – Purkinje cell synapses^33^, excitatory and inhibitory cortical^34,35^ and ribbon synapses^36^. While the distances between VGCCs and the release machinery may vary^37^, a striking parallel is the requirement for very little Ca^2+^ to trigger synaptic vesicle fusion and release^7,38^. Univesicular release at simple lamprey reticulospinal active zones^12^, allows for determination of Ca^2+^ requirements where an AP evokes release of one vesicle at one release site^39^.

Our cell attached recordings indicate a small pool of VGCCs (N, P/Q, R and L-type). Of these, N-, P/Q- and R-type channels contribute to Ca^2+^ signals for central evoked release^40,41^, including in lamprey^9,13,14^. The presence of multiple channel subtypes suggests potential differential occupancy of different slots with varying distance from the primed vesicle^42^ and differential requirements for multiple channel subtypes in evoking neurotransmission. Importantly, even with all channel subtypes available, at each presynaptic terminal, very few Ca^2+^ channels (1-6, mean of 4) from the total pool of channels (mean of 23) open during depolarization. That few channels open during AP evoked depolarization is confirmed by quantitative imaging of the total molar amount of Ca^2+^ that enters each AZ revealing Ca^2+^ entry of ∼4 nMoles per action potential. High speed LLS imaging of single AP-evoked presynaptic Ca^2+^ transients demonstrated that these small numbers of evoked channel opening events are stochastic, both in position within the AZ, and their likelihood of activation upon repeated stimulation. Because the numbers of Ca^2+^ channel opening is low or even absent, these results imply that variance in channel opening may partly explain quantal variation in evoked release of a limited number of primed vesicles paired with few channels.

To further investigate the relationship between primed vesicles and Ca^2+^ channels, we also compared the number of primed vesicles at these terminals with available channels to show a remarkably similar number (23 channels and 22 primed vesicles), precisely enough Ca^2+^ channels available for the number of primed vesicles. There is substantial evidence that primed vesicles form as part of a protein complex that tethers or is tethered by fusion competent Ca^2+^ channels^4,6,23^. Further, entry of small amounts of calcium can be sufficient to trigger release of a single synaptic vesicle^7,23^. Our findings indicate that few channels, perhaps even a single channel, open in extremely close proximity to the primed vesicle, and as in other central terminals^38^, this is sufficient to elevate Ca^2+^ to the hundreds of micromolar concentrations required for Ca^2+^ synaptotagmin interaction^43^ and subsequent fusion^44^. Indeed, prior results in lamprey synapses favor a nanodomain between Ca^2+^ channels and the release machinery, with synaptic transmission being resistant to the slow Ca^2+^ buffer, EGTA^12^. Our present results, thus, imply that primed vesicle complexes each colocalize with a single channel. Of these fusion competent complexes, our data indicate a small number experience channel opening and these openings distribute across the active zone. Thus, we propose that while a single channel is sufficient to evoke fusion, there is a low probability of a channel evoking release. This solution elegantly combines metabolic efficiency of transduction of the AP signal to a fusion event, in which very few Ca^2+^ channels open and only one is required for fusion. The solution implies only a minimal Ca^2+^ load that might otherwise be toxic to be cleared. Nevertheless, the combination of ∼20 primed vesicles with P_r_ of 0.22 allows 100 action potentials to fire before exhausting the primed pool. The RS axons can maintain up to ∼50 Hz firing rates,^45^ leaving 2 s for this exhaustion. This time frame, as in most synapses, presents sufficient time to replenish the primed pool and maintain signaling over long periods. Our findings present a model for efficient synaptic transmission at central synapses that achieves rapid neurotransmission with high temporal precision, while permitting simultaneous maintenance of high transmission fidelity.

## Acknowledgements

We would like to thank Dr. Janet Richmond for helpful discussions and resources throughout the course of this study and Dr. Jonathan Art for feedback on this manuscript. This work was funded by NIH grants R01 MH084874 and R01 NS052699 to SA.

## Methods

All procedures conformed to institutional (Animal Care Committee, University of Illinois at Chicago) and Association for Assessment and Accreditation of Laboratory Animal Care guidelines. Spinal cords were dissected from ammoecoete lampreys (*Petromyzon marinus*), anaesthetized with tricaine methanesulfonate (MS-222, 100 mg/L, Sigma), in ice cold (4°C) Ringer’s solution (mM): 130 NaCl, 2.1 KCl, 2.6 CaCl_2_, 1.8 MgCl_2_, 4 glucose, 5 HEPES, pH 7.6, 270 mOsm.

### Acutely dissociated reticulospinal axon preparation

Briefly, meninges were removed from 1 cm long spinal cord segments, and the dorsal column of the spinal cord removed on a sylgard block in a vibrating slicer in ice-cold Ringer’s solution, leaving intact underneath reticulospinal tracts. The sliced tissue was incubated in a mixture of collagenase and protease in Ringer’s solution (1 mg/ml) at room temperature for 45 minutes. Lateral tracts of the spinal cord were cut and the tissue gently separated on poly-D-lysine (MW>300000, Sigma-Aldrich) coated coverslips by applying mechanical tension along the rostral-caudal axis, until the reticulospinal axons were isolated from the spinal cord interior [Fig.1B]. For details, see Ramachandran *et. al*. 2014^8^.

### Immunohistochemistry

Dissociations were carried out in poly-D-lysine coated petri dishes. Dissociated axons were allowed to recover for 30 minutes at 10°C. The dissociated preparation was fixed in 4% (w/v) paraformaldehyde for 20 minutes, washed with 0.1 M glycine (in PBS), permeabilized with 0.1% Triton-X (in PBS) for 10 minutes and washed with PBS for 20 minutes. All solution changes were performed by perfusion. Blocking was performed with 5% milk (in PBS) overnight at 4°C. Ca^2+^ channels were labeled by incubating with antibodies (host – rabbit; Alomone Labs), specific to N-, P/Q- or R-type VGCCs, (1:200 final equilibrated concentration in dish) for 20 hours at 4°C, labeling a single channel subtype in separate experiments [Fig.S2]. The preparation was washed with PBS for 20 minutes, further blocking carried out with 5% milk (in PBS) for 10 minutes and incubated with secondary antibody Alexa Fluor 633 hydrazide conjugated antibody (host–goat, anti-rabbit; Life Technologies) at a dilution of (1:500 final equilibrated concentration in the dish) for 3 hours at 4°C in dark. Further blocking was carried out with 1% Bovine Serum Albumin (in PBS) for 20 minutes, and washed with PBS for 20 minutes.

Presynaptic AZs were identified by co-labeling carried out by further incubating in Alexa Fluor 488 hydrazide conjugated phalloidin (200 units/ml stock prepared in methanol, final working concentration of 10 units/mL, Life Technologies) for 20 minutes and washed with PBS for 20 minutes. Imaging was carried out on a custom-built upright confocal microscope with a 100X water immersion lens NA 1.0, at 488 and 633 nm. Image processing was carried out using ImageJ. Imaged were post-filtered through a smoothing filter, the two channels were merged and co-localization was determined based on superimposed positioning of VGCC labeling and AZ labeling by phalloidin^46^. Specificity of the antibody labeling was verified by pre-incubating each of the primary antibodies, prior to application, with the corresponding control antigen, for 30 minutes at room temperature.

### Single channel recordings

Dissociations were carried out on poly-D-lysine coated coverslips, in Ringer’s solution, in a custom designed low-noise recording chamber, maintained at 10°C as previously described in Ramachandran *et. al*., 2014^8^. Dissociated axons were allowed to recover for one hour and presynaptic terminals were labeled with FM 1-43 (5 µM) during 30 mM KCl (perfused in Ringer’s solution) evoked stimulation^49^. Excess FM dye was cleared with 1 mM Advasep-7^47^ in Ringer’s solution for 1 minute and washed with Ringer’s solution for 15 minutes. All solutions were applied by perfusion. Single channel recordings were carried out in a stationary bath maintained at 10°C. Imaging was performed using a standard inverted fluorescence microscope fitted with a Hamamatsu Orca CCD Camera and 100X oil immersion lens (NA 1.25). Image acquisition was carried out using Micromanager. A xenon lamp (Nikon) was used as the illumination source and intensity controlled with a neutral density filter to prevent fluorescence bleaching. Patch pipettes (2–10 MΩ) were fabricated from 1.5 mm O.D. thick walled aluminosilicate glass (Sutter Instruments) in a P-87 microelectrode puller (Sutter Instruments), sylgard coated and heat-polished. Low noise (Root mean square (RMS) noise <200 fA) recording conditions were maintained for all recordings. Recordings were only carried out with pipettes where the diameter of the pipette tip was greater than the diameter of the fluorescently labeled target active zone, thus encompassing the entire release face membrane. Mechanical positioning of the patch pipette was carried out using a MP225 Motorized Manipulator (Sutter Instruments). Labeled active zones were patched in the cell-attached configuration to obtain gigaohm seals at 0 mV holding potential and capacitance compensated. ‘n’ reported indicates independent patch recordings, each from a separate dissociated preparation from different animals. The membrane potential of acutely dissociated axons was measured in two ways; by impaling the isolated axon with a microelectrode containing 3 M KCl or by patching the isolated axon in whole cell configuration with a patch pipette containing (mM) 102.5 cesium methane sulphonate, 252 mOsm, pH 7.2. The membrane potential of the acutely dissociated axons was found to be −58 ± 1.6 mV; n = 17 axons), compared to a resting membrane potential of −70 mV in reticulospinal axons in the intact spinal cord. To adjust for this, prior to beginning the step depolarization, the pipette holding potential was adjusted post-seal to +30 mV, which corresponded to a membrane voltage of approximately −80 mV. Step-wise depolarization was evoked using the following stimulation protocol [Fig.2A]: 80 mV pre-pulse for 10 milliseconds (ms) followed 10 ms later by an incremental 10 mV step protocol. A leak step of 10 mV was incorporated for post-acquisition P/10 leak subtraction. RMS noise was monitored in the amplifier and recordings were only carried out if below 200 fA. Data was acquired using an Axopatch 200B (Molecular Devices) with a cooled headstage, in low noise capacitance feedback patch mode, at 50 kHz and filtered using a 5 kHz low-pass 4-pole Bessel filter. VGCC currents were recorded with the following pipette solution: 10 mM CaCl_2_, 80 mM NaCl, 1.8 mM MgCl_2_, 5 mM HEPES, 1 μM Tetrodotoxin (TTX, voltage-gated Na^+^ channels), 20 mM 4-aminopyridine (4-AP, voltage-gated K^+^ channels), 30 mM tetra-ethyl ammonium chloride (TEA-Cl, voltage-gated K^+^ channels), 92 nM Charybdotoxin (ChTx; BK channels), 1 μM Apamin (SK channels), pH 7.6, 270 mOsm. VGCC sub-type specific currents [Fig.2B] were isolated by incorporating subtype specific blockers in the pipette solution, minus the blocker for the desired channel subtype: ω-conotoxin GVIA (N-type, 10 μM); Agatoxin IVa (P/Q-type, 2 μM); SNX-482 (R-type, 2 μM); Nimodipine (L-type, 20 μM). Patch solution for barium recordings [Fig.3B] was: 90 mM BaCl_2_, 5 mM HEPES, 1 μM TTX (voltage-gated Na^+^ channels), 20 mM 4-AP, 30 mM TEA-Cl (voltage-gated K^+^ channels), 92 nM ChTX (BK channels), 1 μM Apamin (SK channels), pH 7.6, 270 mOsm. Cd^2+^ block of VGCC currents was carried out using the following pipette solution: 500 μM CdCl_2_, 10 mM CaCl_2_, 100 mM NaCl, 2.1 mM KCl, 1.8 mM MgCl_2_, 5 mM HEPES, 1 μM TTX, 20 mM 4-AP, 30 mM TEA-Cl, 270 mOsm, pH 7.6. For all data presented, inward currents are plotted with a negative directionality. Detailed analysis of single channel recordings is described in Supplement.

### *In situ* calcium imaging (Molar quantitation of calcium entry)

Dissected 1 cm long spinal cord segments were pinned in a sylgard lined recording chamber and perfused with oxygenated Ringer’s solution, precooled (10 ± 2°C) by passing through a thermocouple Peltier device (Laird). Microelectrodes were fabricated using a P-97 electrode puller (Sutter Instruments) and beveled using a BV-10 Beveler (Sutter Instruments). Reticulospinal axons were impaled with a microelectrode containing fluorescent calcium buffers Fura-2, Oregon-green BAPTA 488 or Fluo5F (Life Technologies) in buffered KCl solution. Once the axon was impaled, the membrane potential was allowed to stabilize to −75 to −80 mV, following which the dye was injected into the axon by applying brief pressure pulses of 35-100 KPa using a picospritzer pressure injector (Parker-Hannifin). Post loading with dye, the axon was stimulated with a train of action potentials, incrementing the frequency of stimulation with each trial (10-100Hz). The axon was allowed to recover between subsequent trials for 3-5 minutes.

For Fura-2, synchronized emission fluorescence measurements were made at 510 nm, exciting at 375 - 385 nm and 350 − 360 nm. Oregon-green BAPTA and Fluo5F imaging was carried out using confocal line scan microscopy. Imaging was carried out with a 40 X (1.1 NA) water-immersion objective and a Hamamatsu Orca CCD camera. At the completion of the calcium imaging, to enumerate the number of presynaptic terminals within the region of measurement, the calcium buffer electrode was withdrawn and the same axon was impaled just distal to the region of measurement with a second electrode filled with Alexa Fluor 594 Hydrazide conjugated Phalloidin (Life Technologies). The phalloidin was pressure injected into the axon and imaging was carried out after 10- 15 minutes allowing for the phalloidin to label presynaptic actin^46^ and a z stack acquired. 3-d reconstruction was carried out post-acquisition and counting of the number of phalloidin labeled puncta was carried out using the ImageJ plugin PointPicker. All fits to the datasets were carried out in Igor Pro (Wavemetrics) and 95% confidence limits of the fits have been displayed [Fig.S4]. See Supplement for detailed description of analysis.

### Lattice light sheet microscopy

Spinal cords were pinned ventral side up in a cooled (10°C) perfused recording chamber under the LLSM objective lenses [Fig.4A]. Spinal reticulospinal axons were prelabeled by pressure injecting the calcium sensitive dye Fluo5F via a sharp micropipette. A stimulating electrode was positioned over the ventro-medial tracts to evoke action potentials in trains (5 at 20 Hz) repeated at 1 min intervals. This evoked Ca transients at active zones (AZs)^9,46^ shown as computed values of ΔF/F after background subtraction [Fig.4B insets] and from which Ca^2+^ diffused from AZs to cause a slower axonal accumulation [Fig.4B]. We imaged Ca^2+^ transients using LLSM at frame rates of up to 800 Hz in single planes across multiple hotspots in a single axon. Transients were imaged in sequential LLSM frames at the same z position. Subtracting the whole axon signal from the AZ signal reveals amplitudes and time courses of the AZ transient [Fig.4Ba-b]. If this subtraction did not show a remaining signal for any stimulation the hotspot was rejected. Additionally, if any stimulation failed to cause Ca^2+^ transients throughout the entire axon, the stimulation was not included in the analysis, as it cannot be determined if the failure was due to a physiological condition or an external failure to generate an action potential. Single event amplitudes were then calculated from exponential fits to the AZ transient decay. See Supplement for detailed description of lattice sheet microscopy and analysis.

### Paired recordings to determine the size of the primed vesicle pool

Paired cell recordings (n = 5) were made between reticulospinal axons recorded with sharp microelectrodes and postsynaptic spinal ventral horn neurons with whole cell patch clamp as previously described^20^. Repeated stimulation of single action potential evoked EPSCs allowed construction of amplitude histograms [Fig.4D] fitted with a multiple gaussians. From this, the number of AZ release sites (n = 7.0 ± 0.5) between each axon/neuron pair, quantal amplitude (*v* = 8.1 ± 2.7 pA) and probability of release at each site (P_r_ = 0.22 ± 0.02) were obtained. To determine the size of the primed vesicle pool, similar paired recordings were made but BoNT/B was included in the presynaptic electrode solution and pressure injected into the axon after obtaining control EPSCs. Light-chain BoNT/B (Calbiochem; 50 μg ml^−1^), was stored at −20°C in 20 mM sodium phosphate, 10 mM NaCl and 5 mM DTT at pH 6.0. The buffered toxins were diluted 2:5 with 2 M potassium methylsulfate and 5 mM HEPES for inclusion in the presynaptic electrode.

### Antibody labeling against Synaptotagmin-1 (Syt-1)

A lumenal domain biotinylated anti-synaptotagmin-1 antibody (rabbit, anti-mouse amino acid residues 1-8, SYSY 105 103BT) diluted 1:50 in Ringer was applied by pressure from a pipette over an isolated axon during 30 mM KCl stimulation of the axon. During subsequent stimulation, streptavidin conjugated dye (Alexa 488) was applied by pipette and fluorescent puncta resolved. Control stimulation and application of streptavidin labeled dye were performed similarly without prior application of antibody against synaptotagmin-1.

