## Supplementary Materials for "Single calcium channel nanodomains drive presynaptic calcium entry at lamprey reticulospinal presynaptic terminals"

Ramachandran, S^1,a^, Rodgriguez S^1^, Potcoava, M^1^. and Alford, S^1,*^.

**Supplementary Material**

**Analysis of single channel recordings**

Leak currents were subtracted and the data filtered through a 1 kHz low pass Gaussian filter in Axograph. Baseline drift adjustment was carried out using the baseline adjustment function in Clampfit.

Determination of single channel current amplitude

Single channel current amplitude, ‘i’, was determined for each VGCC subtype, at indicated voltages, [Fig.2A, Table.S1] from current records visibly demonstrating the opening of only a single channel. Amplitude histograms for the background current noise, measured at holding potential [Fig.2A], were plotted (bin width 50fA) and fitted with a Gaussian function to generate a current noise Gaussian. This Gaussian was scaled to the peak amplitude at zero pA of the current record histogram. It was then subtracted from the channel data amplitude histogram and the subtracted histogram was fitted with a Gaussian [Equation #1]. The single channel current ‘i’ was determined from the centroid of the Gaussian fit. Using this method, ‘i’ was calculated for voltage steps to membrane voltages of -30 mV, 0 mV and 30 mV.

$\mathbf{f}\left( \mathbf{x} \right)\mathbf{=}\mathbf{Ae}^{{\mathbf{-}\left( \frac{\mathbf{(x-C)}}{\mathbf{width}} \right)}^{\mathbf{2}}}$(1)

where C is the centroid, A is the peak amplitude of the Gaussian and $width= \sqrt{2}\sigma$ with $\sigma$ being the standard deviation of the peak.

$\mathbf{f}\left( \mathbf{x} \right)\mathbf{=}{{\mathbf{A}_{\boldsymbol{1}}\mathbf{e}}^{{\mathbf{-}\left( \frac{\mathbf{(x-C)}}{\mathbf{width}} \right)}^{\mathbf{2}}}\mathbf{+}\boldsymbol{A}_{\boldsymbol{2}}\mathbf{e}}^{{\mathbf{-}\left( \frac{\mathbf{(x-(C*2))}}{\mathbf{width}} \right)}^{\mathbf{2}}}$(2)

where A_1_ is the peak amplitude of the first Gaussian, A2 the peak amplitude of the second Gaussian, C the centroid for both the Gaussians and $width= \sqrt{2}\sigma$ with $\sigma$ being the standard deviation of the peak.

Gaussian fits of amplitude histograms

Current from replicate sweeps at a given voltage in a patch recording [referred to as current records, Fig.2A] was binned to generate amplitude histogram plots. These histograms were averaged (between 30-100 sweeps were analyzed) to generate a mean count per sweep amplitude histogram. Bin width for all amplitude histograms was 50 fA. Amplitude histograms were fitted with sum of Gaussian function in Igor Pro.

$\mathbf{f}\left( \mathbf{x} \right)\mathbf{=}{{\mathbf{A}_{\boldsymbol{1}}\mathbf{e}}^{{\mathbf{-}\left( \frac{\mathbf{(x-C)}}{\mathbf{width}} \right)}^{\mathbf{2}}}\mathbf{+}\boldsymbol{A}_{\boldsymbol{2}}\mathbf{e}}^{{\mathbf{-}\left( \frac{\mathbf{(x-(C*2))}}{\mathbf{width}} \right)}^{\mathbf{2}}}\boldsymbol{\ldots\ldots\ldots+}\boldsymbol{A}_{\boldsymbol{n}}e^{{\mathbf{-}\left( \frac{\mathbf{(x-(C*n))}}{\mathbf{width}} \right)}^{\mathbf{2}}}$(3)

where n is the maximum number of channels open (N_chopen.max_), A_1_ is the peak amplitude (peak 0, noise) and C_1_ is the centroid of the noise Gaussian; A_2_ is the peak amplitude at C (centroid), “i*”* (single channel current corresponding to opening of one channel) and A_n_ is the peak amplitude at C times n. n was estimated by dividing the maximum positive amplitude (I_max_) in the histogram demonstrating a bin count (also defined as the peak current observed in m current records from n patches) by “i”; represented by the ratio I_max_/i. Residuals were used to gauge the validity of all fits.

Measurement of tail currents

For each of the VGCC subtypes, tail currents were measured within a window of 20ms, immediately upon repolarization to holding potential of -80 mV [Fig.2A]. Amplitude of individual tail current events were measured and binned (bin width 50 fA) to generate an amplitude histogram, fitted with a sum of Gaussian function [Equation #3]. ‘i’ was determined from the centroid of the Gaussian fit. The number of channels open upon repolarization was determined from the number of Gaussians required to fit the tail current amplitude histogram.

Current – voltage relationships (I-V plots)

For each of the VGCC subtypes, current-voltage (I – V) plots were plotted using the mean single channel amplitude ‘i’ calculated at voltages of -30 mV, 0 mV, 30 mV and upon repolarization to -80 mV (tail current). I –V plots were fitted with a linear function (Equation #4) and the single channel conductance (pS) for each channel subtype was determined by the inverse of the slope. Reversal potential (E_rev_) for each channel subtype was obtained from I–V plots. The X intercept of the linear fit to the I-V data indicates the reversal potential.

$y= a+bx$ (4)

where a is the intercept and b is the slope.

Determination of open probability (P_open_)

Current records were integrated using a trapezoidal algorithm in Igor Pro and open probability was calculated using Equation #5 ^1^.

$P_{open}= \frac{1}{iND}\int_{0}^{D} I\left( t \right)dt$(5)

where P_open_ is the open probability for a single channel at a given voltage, ‘i’ is the single channel current in Amperes, N is number of channels in the current record, D is the duration of the current record and t is time.

**Molar quantitation of Calcium entry**

Spinal cords were isolated and pinned in a custom chamber lined with sylgard and perfused with oxygenated Ringer’s solution (10°C). Sharp microelectrodes were beveled using a BV-10 Beveler (Sutter Instruments) to make it easier to inject dyes into the axon with least possible injection pressure. Reticulospinal axons were impaled with a microelectrode containing the ratiometric dye Fura-2 (Life Technologies) or the non-ratiometric dyes Fluo-5F or Oregon Green BAPTA1 in buffered (5mM HEPES, pH 7.2) KCl solution. Stock solution for dye was prepared in de-ionized water and stored at -20 °C. Dye characteristics (Q and K_d_, see equation 7 for Fura-2 and F_min_, F_max_ and K_d_ see equation 8 for Fluo5F) were calibrated for each stock using Ca^2+^ standards at 10^o^C on the recording microscope. A temperature correction was made for the value of free Ca^2+^ in the standard buffer ^2^.

The dye stock was diluted in buffered KCl (pH 7.2) solution to obtain different final working concentrations of dye. Electrodes were filled through capillary action by placing the shank in the dye solution. Once the axon was impaled, dye was injected into the axon with pressure pulses. To stimulate Ca^2+^ entry, axons were electrically stimulated with a train of action potentials.

For Fura-2 experiments, synchronized emission fluorescence measurements were made at a bandpass of 460-560 nm, while alternating excitation of 375-385 nm and 350-360 nm (Fura-2 isobestic point) using a 40 X (1.1 NA) water-immersion objective and Hamamatsu Orca CCD camera. Absolute dye concentration was calculated by comparing fluorescence of the axon at the isobestic point of Fura-2 with the fluorescence at a similar diameter of the recording electrode in which the dye concentration was known. Fluorescence transient in response to stimulation were obtained at increasing concentrations of dye in the same axon. For Fluo-5F and Oregon Green BAPTA1 measurements, dye was co-injected with Alexa Fluor 594 Hydrazide. Ca^2+^ dye was imaged with excitation at 470 nm and emission at bandpass from 510-570 nm. After each stimulation, the Alexa Fluor 594 Hydrazide was imaged to determine dye concentration in the axon as for Fura-2.

To enumerate the number of presynaptic terminals within the region of measurement, at the completion of the Ca^2+^ imaging, the Ca^2+^ dye containing electrode was withdrawn and the same axon was impaled just distal to the region of measurement with a second electrode backfilled with Alexa Fluor 594 Hydrazide conjugated Phalloidin (Life Technologies). Phalloidin was pressure injected into the axon and imaging was carried out after 10-15 minutes allowing for the phalloidin to label presynaptic actin^3^. The axonal volume was reconstructed from acquired series of z slices. The number of phalloidin puncta was counted in these images. The mean number of AZs per axon region imaged was 79 ± 7. All fits to the datasets [Fig.S4] were carried out in Igor Pro and 95% confidence limits of the fits displayed.

Calcium dye was used as a buffer to quantify amounts of evoked Ca^2+^ entry to lamprey axons. To achieve this buffer capacities of dye injected into the axons and known concentrations were calculated. Properties of the Ca^2+^ transient were determined over a range of buffering capacities of the dye by the use of dyes with 3 different affinities over a range of concentrations. The buffering capacity during each stimulated transient was calculated using equation 6 ^4–6^.

$\kappa_{dye}=\frac{\Delta\left[ {Ca}_{dye} \right]}{{\Delta\left[ {Ca}^{2+} \right]}_{i}}=\frac{\left[ {Dye}_{total} \right]}{k_{d}\left[ \left( 1+\frac{\left[ {Ca}^{2+} \right]_{2}}{k_{d}} \right)\left( 1+\frac{\left[ {Ca}^{2+} \right]_{1}}{k_{d}} \right) \right]}$	(6)

where [Ca_dye_] is the concentration of Ca^2+^-bound dye, Δ[Ca^2+^]_i_ is the evoked change in Ca^2+^ concentration, and [Dye_total_] is the total dye concentration. [Ca^2+^]_1_ and [Ca^2+^]_2_ are the resting Ca^2+^ and peak Ca^2+^ concentrations before and during stimulation respectively. They are derived from the equation for 1:1 complexation of Ca^2+^ and dye as a buffer. For ratiometric measurements using Fura 2 can be expressed as in equation 7^7^ and for non-ratiometric dyes as equation 8.

$\left[ {Ca}^{2+} \right]=K_{d}Q\frac{\left( R-R_{min} \right)}{\left( R_{max}-R \right)}$	(7)

and for non-ratiometric dyes $\left[ {Ca}^{2+} \right]_{i}=K_{d}\frac{\left( F-F_{min} \right)}{\left( F_{max}-F \right)}$	(8)

Where R is the ratio of fluorescence intensities at 350-360 nm (Fura-2 isobestic point) and 380 nm excitation, Q is the ratio of fluorescence minimum to fluorescence maximum at 380 nm for ratiometric dyes. F_max_ for each transient was determined by intense stimulation to saturation after the transient was measured. F_min_ was calculated as a ratio of this value from measurement of Ca^2+^ standards. Values of F calculated throughout the transient as ∆F/(F+1) could then be used to calculate [Ca^2+^]_i_ from equation 8.

Using these data, the endogenous buffering capacity of the axon (κ_end_) was determined from the dependency of the rate of decay the dye recorded Ca^2+^ transient to the buffering capacity of the dye. The recovery rate of the stimulus-coupled fluorescence change was fitted with an exponential function and the decay rate (τ) measured, over a range of κ_dye_. These values of τ were plotted against κ_dye_ and this data was fitted with a linear function [Fig.S4] yielding an estimate of the endogenous buffering capacity (κ_end_) to be 13.1 ± 11.1 from equation 9, which describes the rate (τ) of a pulse-like Ca^2+^ signal in a cell compartment.

$\tau=\tau_{ext}(1+\kappa_{end}+\kappa_{dye})$	(9)

This value of κ_end_ gives an estimate independent of the absolute measure of Ca^2+^ transient amplitudes and thus provides a reasonable measure of validity for the remaining calculations.

We may also calculate the free Ca^2+^ concentration evoked by stimulation and the total molar quantity of Ca^2+^ that enters the axon. By measuring the peak amplitude of the free Ca^2+^ transient throughout the varicosity over a range of values of κ_dye_^6,8^, we may state:

$\Delta\left[ {Ca}^{2+} \right]_{i}=\frac{\Delta\left[ Ca \right]_{total}}{\left( 1+\kappa_{dye}+\kappa_{end} \right)}$	(10)

Where ∆[Ca^2+^]_i_ varies with κ_dye_. Values of ∆[Ca^2+^]_i_ are computed from our data using equations (7 or 8) and values of κ_dye_ from equation (6). For each stimulus, as the value of κ_dye_ rises it will come to dominate binding of Ca^2+^ entering the cell compartment. Thus, the value of ∆[Ca_dye_] can be calculated for each stimulus as the value of κ_dye_ increases.

$\Delta\left[ Ca \right]_{total}=\Delta\left[ {Ca}_{dye} \right].\left\{ \frac{\left( \kappa_{end}+1 \right)}{\kappa_{dye}}+1 \right\}$	(11)

Values of ∆[Ca]_total_, and κ_end_ can be determined by extracting constants from fits of either equation (10) or (11). The true value of peak ∆[Ca^2+^]_i_ in the axons when no dye is present is obtained by extrapolating the fit to the y intercept in equation (10) where κ_dye_ = 0. This is the Ca^2+^ concentration measured in the entire volume of the axon. From this we can calculate the total Ca^2+^ entering the axons using either equation (10) or (11). We also calculated axon volumes from images assuming the axons are cylindrical. The mean molar amount of Ca^2+^ entering at each terminal can then be given by dividing this total by the number of synaptic terminals imaged. These were obtained from counting phalloidin clusters^3,9^ from the same axons at the end of the experiment.

Peak Δ[Ca^2+^], in the volume of axon visualized, was plotted for a range of calculated κ_dye_ and fitted with equation 10 yielding a peak free ∆[Ca^2+^] of 31.6 ± 5.2 nM throughout the axon per stimulus from the intercept on the ordinate axis [Fig.S4]. This fit also yielded the value of total Ca^2+^ concentration, bound and unbound (∆[Ca]_total_) of 109.9 ± 19.2 nM that entered the whole axon volume and a κ_end_ of 2.5 ± 1.1. Finally, a plot of ∆[Ca]_total_ for each stimulus against k_dye_, fitted by equation 11 yields a κ_end_ of 5.2 ± 2.2, an asymptotic value of ∆[Ca]_total_ representing full dye binding of all Ca^2+^ entry of 116.9 ± 13.2 nM. These approaches yielded similar values of each of these parameters with mean values listed in Table.S6 from which total charge entering and charge per AZ could be calculated. Note that during Ca^2+^ entry that occurs only at AZs, the local concentration of Ca^2+^ is transiently at much higher concentrations which evokes neurotransmission, but this high concentration almost immediately disperses throughout the much larger axonal volume from which it was imaged. From these calculations [Table.S6] we conclude a total charge entering the axon at each AZ during one AP is just 7.98 ± 0.90 fC which represents a current of 1.6 ± 0.3 pA over the duration of the action potential (2 to 3 ms of depolarization and 2 ms of estimated tail current). Mean channel currents over this voltage range were approximately 0.4 pA (from the integral of slopes between -80 and +20 mV in Fig 2C), indicating opening of a mean of 4 channels per active zone per AP. This is very similar to results from cell attached recordings and quantal imaging of Ca^2+^ transients providing independent validation of those approaches.

**Fluorescence Analysis of Ca^2+^ entry at AZs with LLSM**

To analyze variance of presynaptic Ca^2+^ entry through few channels we must resolve Ca^2+^ responses, spatially and temporally at individual AZs, evoked by single action potentials. Wide field fluorescence causes high fluorescence background noise and rapid photobleaching from excitation light. Typically, this background noise has been avoided by confocal or two photon approaches, but high excitation intensities cause rapid photo-bleaching that limits the number of repeat scans achievable. Point scanning approaches limit the temporal resolution of recording, and expanding this to line-scanning still limits the field of view. This, and the limited number of repeated exposures caused by photobleaching, make interpretations of variations in responses between active zones or within single active zones not feasible.

To overcome these limitations, we built a custom Lattice Light Sheet Microscope (LLSM)^10^. The LLSM generates a convergent lattice of 30 Bessel beams^10,11^, confined to a plane (~0.4 µm deep) as they self-reinforce and propagate well in situ, providing substantially better resolution than conventional light sheet imaging. The LLSM beam is viewed orthogonally to its projection allowing low excitation intensity illumination of planes to be viewed rapidly in sequence with little photobleaching, allowing many repeated measurements of action potential evoked transients. The self-reinforcing nature of the Bessel beam provides improvements of light sheet penetration^10^. This provides a volumetric imaging approach, but in which we image an entire plane at high temporal resolution (up to 800 Hz). We can record Ca^2+^ transients at multiple active zones at once, without significant background noise and photobleaching, and when we capture more than one active zone, we can confirm triggering of an AP by analysis of multiple hotspots.

Repeated stimulation and recording of Ca^2+^ hotspots using LLSM enabled high signal to noise transients to be repeatedly recorded. Peak amplitudes of values of ∆F/F were calculated and plotted as histograms. Multiple gaussian fits were applied to this using equation 12.

$f\left( F \right)=A_{noise}.e^{\left\{ {-\left( \frac{F-F_{noise}}{W_{noise}} \right)}^{2} \right\}}+A_{1}.e^{\left\{ {-\left( \frac{F-F_{t}}{W_{t}} \right)}^{2} \right\}}+A_{2}.e^{\left\{ {-\left( \frac{F-{3.F}_{t}}{{3.W}_{t}} \right)}^{2} \right\}}......+A_{n}.e^{\left\{ {-\left( \frac{F-{n.F}_{t}}{{n.W}_{t}} \right)}^{2} \right\}}$ (12)

Where *A_noise_* = peak number of failures at F*_noise_* – value of ΔF/F of failure; *A_1_, A_2_* to *A_n_* = peak numbers of event at *F_t_* – the transient peak ΔF/F, and its multiples. *W_t_* = variance of the unitary amplitude.

**Supplementary Figure 1**

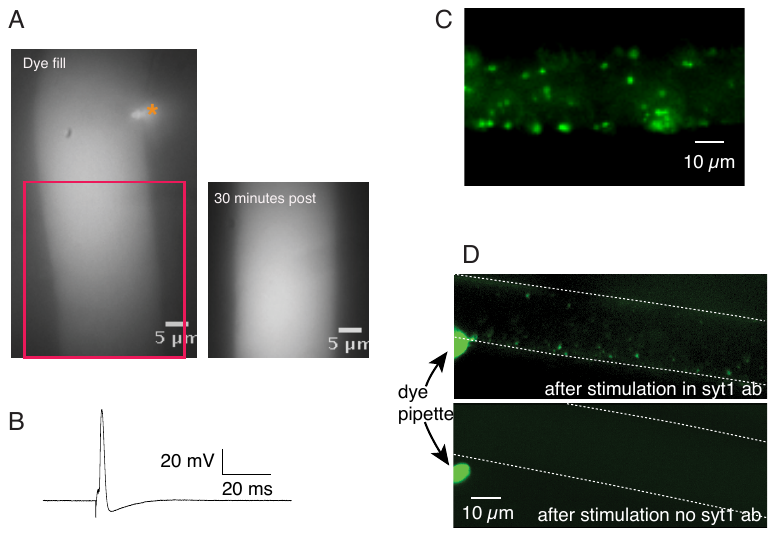

A. Representative images showing Alexa Fluor 488 Hydrazide filling of dissociated reticulospinal axon (left) by pressure injection of the dye through a patch pipette (orange asterix) in whole cell patch configuration. Note 30 minutes later, the dye integrity is maintained within the axon (red outlined region of the axon in left panel upon dye loading, right panel post 30 minutes), indicating that the axons remain structurally intact during and after dissociation.
B. Representative action potential (n=7 axons) elicited in a dissociated reticulospinal axon by stimulating with a sharp microelectrode containing 3M KCl.
C. Representative image of a region in a dissociated reticulospinal axon showing punctate FM 1-43 labeled recycling vesicle clusters, labeled by 30 mM KCl bath-perfusion stimulation. Excess FM1-43 was removed with Advasep-7^12^.
D. Antibody labeling against Synaptotagmin-1 (Syt-1). A lumenal domain biotinylated anti-synaptotagmin 1 antibody (rabbit, antimouse amino acid residues 1-8, SYSY 105 103BT) diluted 1:50 in Ringer was applied by pressure from a pipette over an isolated axon during 30 mM KCl stimulation of the axon. During subsequent stimulation, streptavidin conjugated dye (Alexa Fluor 488 Hydrazide) was applied by pipette. Fluorescent puncta were again resolved. Control stimulation and application of streptavidin labeled dye, but without prior application of antibody against synaptotagmin-1 revealed no puncta.

**Supplementary Figure 2**

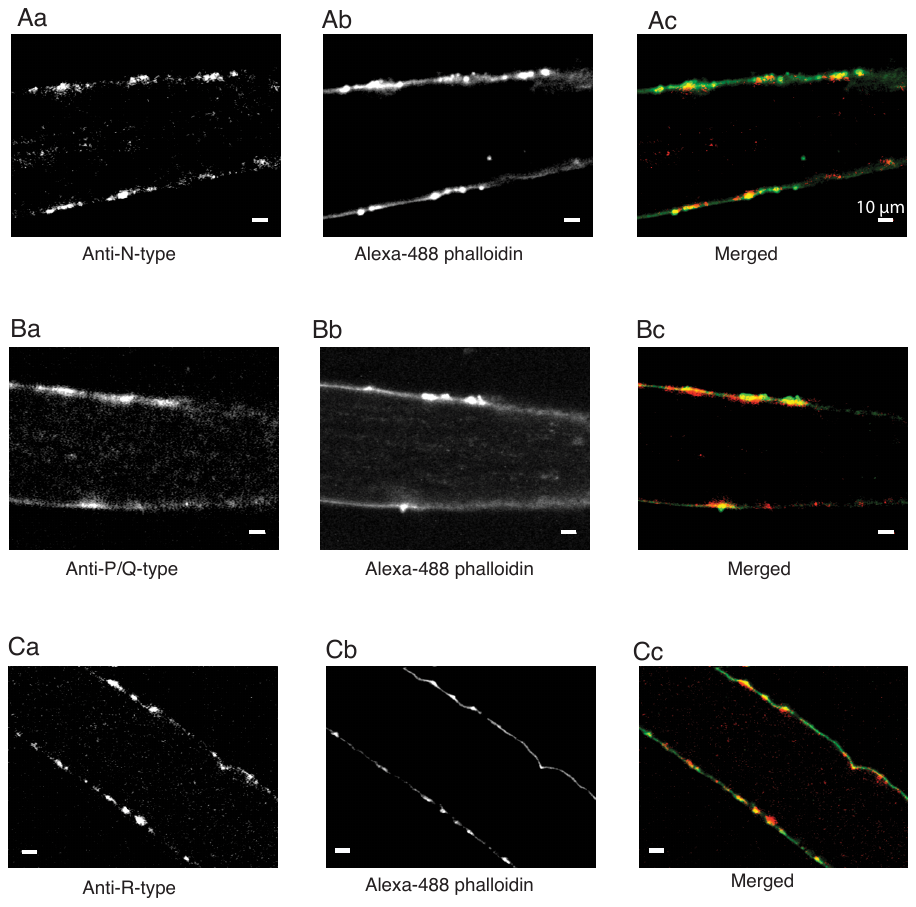

Antibody labeling against N-type (CaV2.2) [Aa], P/Q-type (CaV2.1) [Ba] and R-type (CaV2.3) VGCC [Ca] demonstrating punctate distribution along the axon membrane. Note: Primary antibodies against VGCC subtypes (rabbit, Alomone Labs); secondary antibody Alexa Fluor 633 Hydrazide conjugated (goat, anti-rabbit, Alomone Labs). The primary anti N-type antibody was against an intracellular epitope (C)RHHRHRDRDKTSASTPA, corresponding to amino acid residues 851- 867 of rat CaV 2.2 (Accession Q02294) located in the intracellular loop between domains II and III of the VGCC. The primary anti-P/Q-type antibody was against an intracellular epitope (C)PSSPERAPGREGPYGRE, corresponding to amino acid residues 865-881 of rat CaV 2.1 (Accession P54282) located in the intracellular loop between domains II and III of the VGCC. The primary anti-R-type antibody was against an intracellular epitope (C)SASQERSLDEGVSIDG, corresponding to amino acid residues 892-907 of rat CaV 2.3 (Accession Q07652) located in the intracellular loop between domains II and III of the VGCC. Presynaptic terminal locations were determined in the same axon by labeling presynaptic actin with Alexa Fluor 488 Hydrazide conjugated phalloidin [Ab, Bb, Cb]. Colocalization of VGCC labeling with locations of presynaptic terminals for each VGCC subtype characterized Red indicates VGCC labeling, green indicates presynaptic actin labeling and yellow colocalization. [Ac, Bc, Cc]. Scale bar in all images 10 µm. n for each VGCC subtype: N-type 5 axons from 3 animals; P/Q-type 9 axons from 4 animals; R-type 3 axons from 2 animals.

**Supplementary Figure 3**

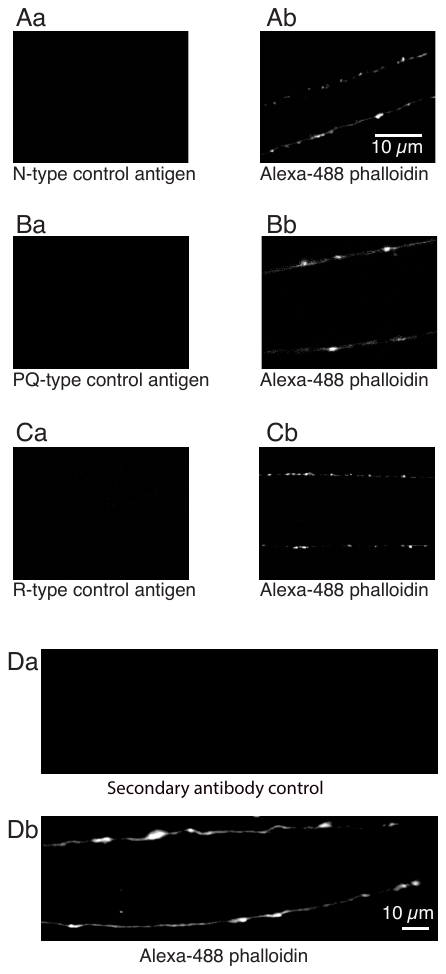

[A-C] Representative images showing controls for the primary antibodies used for N-type, P/Q-type and R-type channel staining. Primary antibody was preincubated with specific control antigen for 30 minutes before staining the preparation [Aa-Ca]. Note the absence of any VGCC labeling in the controls.

[Ab-Cb] show the presynaptic terminals location labeled by Alexa Fluor 488 Hydrazide conjugated phalloidin labeling of presynaptic actin in same region of the axon as in the left panel. Scale bar in all images 10 μm. n for each control: N-type control antigen 10 axons from 2 animals; P/Q-type control antigen 5 axons from 2 animals; R-type control antigen 3 axons from 2 animals

[D] Representative image of control for secondary antibody [Da] and presynaptic terminals location labeled by Alexa Fluor 488 Hydrazide conjugated phalloidin labeling of presynaptic actin [Db] in same region of the axon. n = 8 axons from 3 animals.

**Supplementary Figure 4**

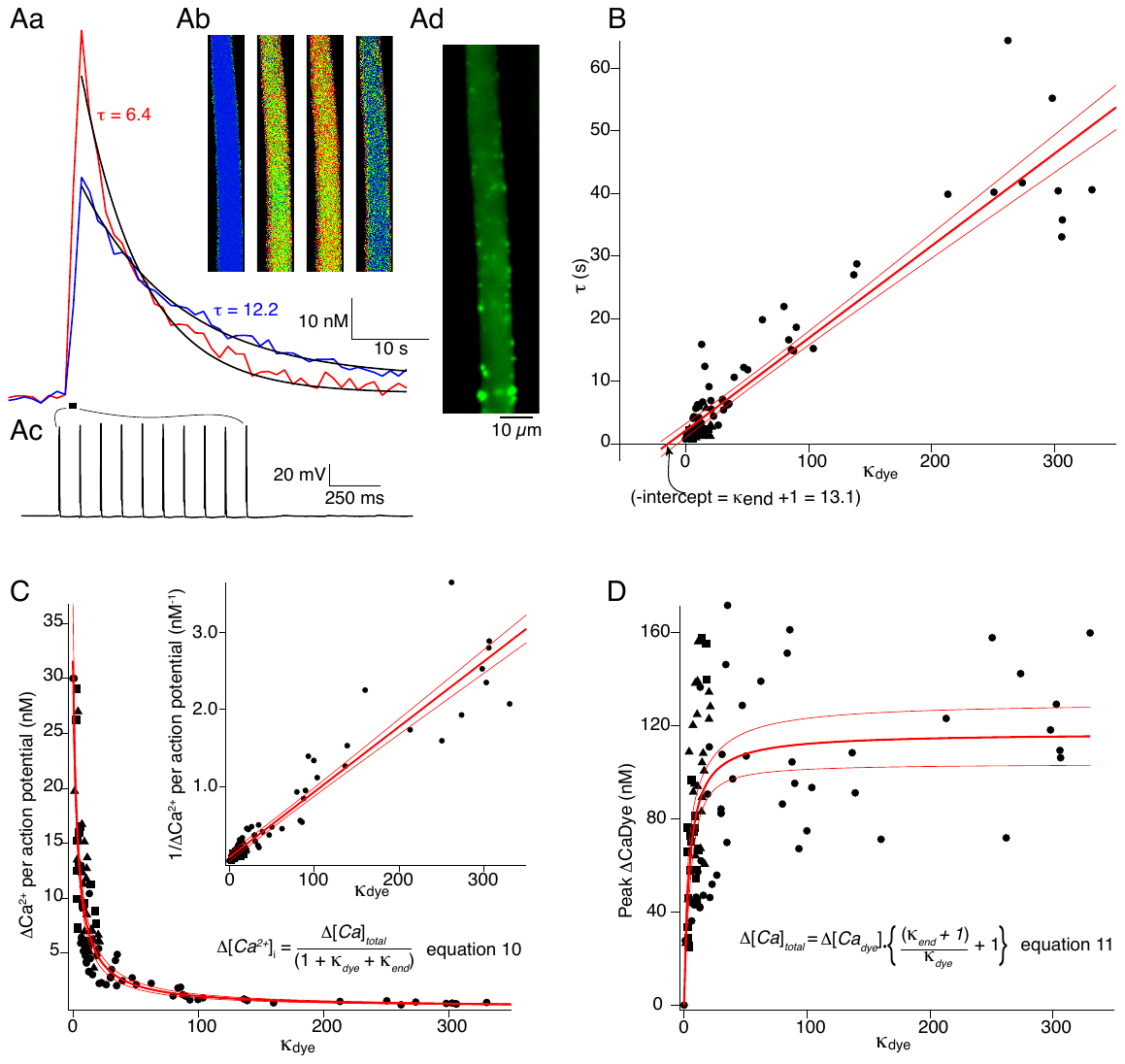

To address whether acute dissociations disrupted VGCC numbers at individual presynaptic active zones, we verified this number using quantitative Ca^2+^ imaging in which stimulus evoked Ca^2+^ influx into the presynaptic terminal was characterized by introducing exogenous fluorescent calcium buffers.

A. [Aa-Ab] Reticulospinal axons were impaled with a microelectrode containing the ratiometric Ca^2+^ dye Fura-2 (Life Technologies) or the non-ratiometric dyes Fluo-5F or Oregon Green BAPTA1 co-injected with an inert dye to calculate injected concentration. Dyes were in buffered KCl solution (pH 7.2). Dye was injected into the axon with pressure pulses. Axons were stimulated with a sequence of action potential trains [Ac] between which more dye was pressure injected to obtain responses at a range of dye concentrations and therefore k_dye_. Representative Ca^2+^ responses showing the measured decay rate (τ). [Ad] At the completion of the Ca^2+^ imaging, to enumerate the number of presynaptic terminals within the region of measurement, the Ca^2+^ dye containing electrode was withdrawn and the same axon was impaled just distal to the region of measurement with a second electrode backfilled with Alexa Fluor 594 Hydrazide conjugated Phalloidin (Life Technologies). Phalloidin was pressure injected into the axon and imaging was carried out after 10-15 minutes allowing for the phalloidin to label presynaptic actin surrounding vesicle clusters identifying presynaptic terminal locations.
B. Endogenous buffering capacity of the axon (κ_end_) was determined from the dependency of the rate of decay the dye recorded Ca^2+^ transient to the buffering capacity of the dye. The recovery rate of the stimulus-coupled fluorescence change was fitted with an exponential function and the decay rate (τ) measured [Aa]. τ was measured over a range of κ_dye_. values of τ were plotted against κ_dye_ and this data was fitted with a linear function (dark red line, 95% confidence indicated by encompassing faint red lines) yielding an estimate of the endogenous buffering capacity (κ_end_) to be 13.1 ± 11.1 from equation (9).

**C-D.**

Peak Δ[Ca^2+^], in the region of measurement, was plotted for a range of calculated k_dye_ and fitted (dark red line, 95% confidence indicated by encompassing faint red lines) with equation (13) yielding a peak ∆[Ca^2+^] of 31.6 ± 5.2 nM per stimulus from the intercept on the ordinate axis. This fit also yielded value of total Ca^2+^ concentration, bound and unbound (∆[Ca]_total_) of 109.9 ± 19.2 nM and a κ_end_ of 2.5 ± 1.1. A plot of ∆[Ca]_total_ for each stimulus against k_dye_, fitted by equation 14 yields a κ_end_ of 5.2 ± 2.2, an asymptotic value of ∆[Ca]_total_ representing full dye binding of all Ca entry of 116.9 ± 13.2 nM. These approaches yielded similar values of each of these parameters with mean values listed in Table.S6 from which total charge entering and charge per AZ could be calculated. From these calculations (Table.S6) we conclude a total charge entering the axon at each AZ during one AP is 11.2 fC which represents a current of 2.79 pA over 5 ms (the rise time of the response). This is very similar to results from cell attached recordings.

**Supplementary Table 1**

| Voltage (mV) | -80 (tail current) | -30 | 0 | 30 |  |
| --- | --- | --- | --- | --- | --- |
|  | i (pA) | i (pA) | i (pA) | i (pA) | pS |
| **N-type** | 0.58 ± 0.01 | 0.35 ± 0.03 | 0.28 ± 0.01 | 0.26 ± 0.01 | 2.97 ± 0.62 |
| **PQ-type** | 0.47 ± 0.01 | 0.34 ± 0.01 | 0.29 ± 0.03 | 0.27 ± 0.02 | 1.82 ± 0.36 |
| **R-type** | 0.47 ± 0.01 | 0.32 ± 0.02 | 0.28 ± 0.02 | 0.26 ± 0.01 | 1.94 ± 0.41 |
| **L-type** | 0.53 ± 0.003 | 0.31 ± 0.01 | 0.28 ± 0.02 | 0.27 ± 0.01 | 2.4 ± 0.75 |

Mean single channel current measured for each VGCC subtype. Single channel conductance (pS) for each VGCC subtype was obtained from the slope of current – voltage relationship plots (I-V plots). Data represented as mean ± SE. n (patch recordings) for each channel subtype: N-type (n=6), P/Q-type (n=5), R-type (n=6), L-type (n=4)

**Supplementary Table 2**

| mean P_open_ |  |  |  |
| --- | --- | --- | --- |
| Voltage (mV) | -30 | 0 | 30 |
| N-type | 0.17 ± 0.04 | 0.26 ± 0.03 | 0.24 ± 0.02 |
| P/Q-type) | 0.19 ± 0.04 | 0.29 ± 0.03 | 0.24 ± 0.01 |
| R-type | 0.14 ± 0.04 | 0.22 ± 0.02 | 0.18 ± 0.01 |
| L-type | 0.22 ± 0.05 | 0.24 ± 0.02 | 0.18 ± 0.02 |

Mean open probability (P_open_) calculated for each VGCC subtype at -30, 0 and 30 mV. P_open_ was calculated by integrating current data from current records (Equation #5), at each analyzed voltage (-30, 0 and 30 mV), clearly demonstrating the opening of only a single channel. Data reported as mean ± SE. n (patch recordings) for each channel subtype: N-type (n=6), P/Q-type (n=5), R-type (n=6), L-type (n=4)

**Supplementary Table 3**

Maximum number of VGCCs observed to open at individual presynaptic active zones

| **Voltage (mV)** | **-30** | **0** | **30** |  |
| --- | --- | --- | --- | --- |
| **N-type** |  |  |  |  |
| **I_max_ (pA)** | 1.6 | 1.7 | 2.7 |  |
| **i (pA)** | 0.35 | 0.28 | 0.26 |  |
| **I_max_/i** | 4.57 | 6.07 | 10.4 |  |
| **Nch_open.max_** | 4 | 6 | 10 |  |
| **Nch_patch.max_** |  |  |  | 10 |
| **P/Q-type** |  |  |  |  |
| **I_max_ (pA)** | 1.35 | 2.85 | 2.15 |  |
| **i (pA)** | 0.34 | 0.29 | 0.27 |  |
| **I_max_/i** | 3.97 | 9.83 | 7.96 |  |
| **Nch_open.max_** | 3 | 9 | 7 |  |
| **Nch_patch.max_** |  |  |  | 9 |
| **R-type** |  |  |  |  |
| **I_max_ (pA)** | 1.3 | 5.35 | 8.35 |  |
| **i (pA)** | 0.32 | 0.28 | 0.26 |  |
| **I_max_/i** | 4.06 | 19.11 | 32.12 |  |
| **Nch_open.max_** | 4 | 19 | 32 |  |
| **Nch_patch.max_** |  |  |  | 32 |
| **L-type** |  |  |  |  |
| **I_max_ (pA)** | 3.35 | 2.6 | 4.8 |  |
| **i (pA)** | 0.31 | 0.28 | 0.27 |  |
| **I_max_/i** | 10.81 | 9.29 | 17.78 |  |
| **Nch_open.max_** | 10 | 9 | 17 |  |
| **Nch_patch.max_** |  |  |  | 17 |
| **Total Nch_patch.max_**  **(N+P/Q+R+L)** |  |  |  | 68 |
| **Total Nch_patch.max_ (N+P/Q+R)** |  |  |  | 51 |

Channel numbers enumerated from patch recordings at single presynaptic AZs, with10 mM [Ca^2+^]_external_ in the patch pipette, under conditions where currents specific to a single VGCC subtype were pharmacologically isolated. For each indicated channel type, measurements were consolidated from all current records in all obtained patch recordings. [Fig.2B,3A]

I_max_ (pA) - Maximum calcium current amplitude recorded. I_max_ at each of the voltages is the peak current in m current records from n patches or can also be read from the mean count/ maximum current value at which a bin count could be observed in the mean count/current record amplitude histogram.

i (pA) - single channel amplitude

N_chopen.max_ for a given voltage is the maximum number of channels simultaneously opening in n patches.

N_chpatch.max_ is the maximum number of channels simultaneously opening at a presynaptic terminal for each VGCC subtype.

n (patch recordings) for each channel subtype: N-type (n=6), P/Q-type (n=5), R-type (n=6), L-type (n=4)

**Supplementary Table 4**

Number of VGCCs at individual presynaptic active zones

|  | **N_chpatch_** | **N_chpatch.mean_** |
| --- | --- | --- |
| **N -type** | 4 - 10 | 5 |
| **P/Q - type** | 3 - 9 | 6 |
| **R - type** | 4 - 32 | 12 |
| **L - type** | 3 - 17 | 10 |
| **Total N_chpatch.mean (N+P/Q+R+L)_** |  | 33 |
| **Total N_chpatch.mean (N+P/Q+R)_** |  | 23 |

Channel numbers enumerated from patch recordings at single presynaptic AZs with 10 mM [Ca^2+^]_external_ in the patch pipette under conditions where currents specific to a single VGCC subtype were pharmacologically isolated. For each indicated channel type, measurements were consolidated from all current records in all obtained patch recordings. [Fig.2B,3A]

N_chpatch_ is the number of channels observed at a presynaptic terminal, reported as a range (minimum-maximum).

N_chpatch.mean_ is the mean number of channels at a presynaptic terminal.

n (patch recordings) for each channel subtype: N-type (n=6), P/Q-type (n=5), R-type (n=6), L-type (n=4)

**Supplementary Table 5**

| **Voltage (mV)** | -30 | 0 | 30 |  |
| --- | --- | --- | --- | --- |
| **i (pA)** | 0.4 | 0.35 | 0.3 |  |
| **I_max_ (pA)** | 0.9 | 1.4 | 1.65 |  |
| **I_max_/i** | 2.3 | 4 | 5.5 |  |
| **N_chopen.max_** | 2 | 4 | 6 |  |
| **N_ch.patch_** |  |  |  | 2-6 |
| **N_ch.patch.mean_** |  |  |  | 4 |

Number of channels opening at a presynaptic terminal, in response to a depolarizing stimulus [Fig.2A], under conditions where all VGCCs are available for opening. Channel numbers enumerated from patch recordings (n=5) at single presynaptic AZs with 90 mM [Ba^2+^]_external_ in the patch pipette. For each indicated channel type, measurements were consolidated from all current records in all obtained patch recordings. ‘i’ (pA) is the single channel amplitude at each of the voltages and can be estimated as the centroid of the sum of Gaussian fits of the amplitude histogram [Fig.3B]. I_max_ at each of the voltages is the peak current observed from n patches. N_chopen.max_ for a given voltage is the maximum number of channels opening observed in n patches. N_chpatch_ is the number of channels opening observed at a presynaptic terminal, reported as minimum-maximum, and N_chpatch.mean_ is the mean number of channels opening at a presynaptic terminal from n patches.

**Supplementary Table 6**

Quantitation of molar calcium entry into axons and active zones

| Parameter | Value | Source |
| --- | --- | --- |
| κ_end_ | 6.9 | Mean from fits to Fig S4 |
| Resting free [Ca^2+^]_i_ (nM) | 150 ± 21 | Mean of Fura 2 measurements |
| Peak free Δ[Ca^2+^]_i_ / stimulus (nM) | 36.8 ± 5.2 | Fig S4C |
| Number of presynaptic terminals | 79 ± 7 | Phalloidin labeling |
| Mean axon volume µm^3^ | 28100 ± 2200 | Diameter of axons |
| Total axonal [Ca^2+^] (Δ[Ca_total_] / Stimulus) (nM) | 116.9 ± 13.2 | Fig S4D |
| Total molar quantity Ca^2+^ entering (Moles)  Total # ions entering  Total charge (C)  Current over 5 ms (pA) | 4.14x10^-9^ ± 0.46 x10^-9^  24900 ± 2800  7.98 x 10^-15^ ± 0.90 x10^-15^  1.60 ± 0.23 | calculated |

Table summarizes the quantification of action potential-evoked entry of calcium into presynaptic axons and individual active zones. This enables an estimate to be made of the current carried by calcium at each action potential.

**Supplementary table 7: Quantal release of neurotransmitter**

| Parameter | Value | Source |
| --- | --- | --- |
| Number of AZ release sites between pairs of axons and target neurons | 7 ± 0.5 |  |
| Quantal amplitude (pA) | 8.1 ± 2.7 | Fig.4 |
| Probability of release at each release site | 0.22 ± 0.02 | Fig.4 |

Quantal analysis of evoked responses from paired recordings between individual axons and their target neurons enabled calculation of probability of release at each active zone.
